# Tunable PhenoCycler Imaging of the Murine Pre-Clinical Tumour Microenvironments

**DOI:** 10.1101/2023.09.18.558299

**Authors:** Madelyn J. Abraham, Christophe Goncalves, Paige McCallum, Vrinda Gupta, Samuel E. J. Preston, Fan Huang, Hsiang Chou, Natascha Gagnon, Nathalie A. Johnson, Wilson H. Miller, Koren K. Mann, Sonia V. del Rincon

**Author notes:** Correspondence to: Sonia V. del Rincon; Koren Mann; Wilson H. Miller Jr.

## Abstract

The tumour microenvironment (TME) consists of tumour-supportive immune cells, endothelial cells, and fibroblasts. PhenoCycler, a high-plex single cell imaging platform, is used to characterize the complexity of the TME. Here, we used PhenoCycler to spatially resolve the TME of 8 routinely employed pre-clinical models of lymphoma, breast cancer, and melanoma. Our data reveal distinct TMEs in the different cancer models that were imaged, and show that cell-cell contacts differ depending on the tumour type examined. For instance, we found that the immune infiltration in a murine model of melanoma is altered in cellular organization in melanomas that become resistant to αPD-1 therapy, with depletions in a number of cell-cell interactions. Furthermore, we provide detailed pipelines for the conjugation of antibodies that are optimized for PhenoCycler staining of murine FFPE tissues specifically, alongside open-source data analysis procedures. Overall, this is a valuable resource study seamlessly adaptable to any field of research involving murine models.

## Introduction

Over the past two decades, there has been growing appreciation for the role of the tumour microenvironment (TME) in cancer biology (1, 2). As such, the central dogma of tumour progression has evolved to assert that oncogenic mutations underlie the transformation of normal cells to malignant cells, and subsequently, non-transformed cells are recruited via secretion of soluble factors, such as cytokines, chemokines, and extracellular vesicles, to support further cancer cell survival and propagation (3–6). The non-transformed cellular elements of the TME, including immune cells, fibroblasts, and endothelial cells, interact with tumour cells, and both cellular composition and intercellular interactions within the TME are critical influencers of cancer cell growth, metastasis, and response to therapy. Many emerging therapeutics, most notably immune checkpoint inhibitors (ICIs), specifically target components of the TME to elicit tumour control.

Phenotyping of the murine TME has helped to understand the response to novel combinatorial therapies and to track changes in tumour progression from initiation to metastatic disease (7, 8), with multi-parameter flow cytometry being the most widely used technique to study the composition of the TME (9). In this method, malignant tissues are dissociated into single cell suspensions, stained with a panel of antibodies, and run through a flow cytometer, allowing for the identification of cells within the TME. However, a recent body of work has highlighted that TME composition alone is only part of a much bigger picture, and spatial information (e.g. cell-cell interactions) is crucial to further understand tumour progression and response to treatment. Immunofluorescence (IF) imaging of tumour sections, on the other hand, can preserve tissue architecture but is usually restricted to detection of 1 or 2 markers. To overcome these limitations, a surge of highly multiplexed tissue imaging technologies has emerged in the last 10 years (10–13), aimed at providing single cell spatial phenotyping of the TME and other complex tissue types.

PhenoCycler, formerly known as CODEX (CO-Detection by indexing (13)), has shown immense promise in the highly multiplexed imaging space. In brief, antibodies targeting desired proteins are conjugated to unique oligonucleotide “barcodes” and are then used to stain fresh frozen or formalin-fixed paraffin-embedded (FFPE) tissues. The PhenoCycler instrument is then used to automate the cyclic process of tissue washing, hybridizing up to three fluorescent “reporters” to primary antibodies oligonucleotide “barcodes”, imaging the tissue, then removing the fluorescent reporters before starting a new cycle process. This iterative process is repeated until all antibodies in a staining panel have been visualized (14). Reporters are complementary oligonucleotides to the unique barcodes, and are tagged with either fluorophores ATTO550 AF647, or AF750. As of this writing, PhenoCycler has been used to image up to 101 different markers in single tissue (15, 16), and has been used to spatially profile human cancers such as cutaneous T cell lymphoma (17), follicular lymphoma (18), diffuse large B cell lymphoma (19), Hodgkin’s lymphoma (20), bladder cancer (21), colorectal cancer (22), basal cell carcinoma (23), glioblastoma (24), breast cancer (25), and head and neck squamous cell carcinoma (26), and human non-cancerous conditions such as ulcerative colitis (27), diabetic nephropathy (28), functional dyspepsia (29), vitiligo (30), and Alzheimer’s disease (31).

Comparatively fewer publications have used PhenoCycler technology to image murine tissues, and all have reported staining for fresh-frozen samples (13, 32–36). However, many research groups maintain archives of FFPE murine tissues. FFPE tissue blocks can be successfully sectioned and imaged with minimal evidence of degradation for up to 30 years (37), and FFPE tissues from multiple cohorts or experimental conditions can be easily combined into a single tissue microarray (TMA). Thus, we aimed to develop a tunable murine PhenoCycler antibody panel optimized for FFPE staining, thereby enabling researchers to utilize their archival materials to test newly developed hypotheses with existing material and bypassing the need to perform new mouse studies.

Herein we show TME data obtained using 16-plex PhenoCycler staining on FFPE tissues from pre-clinical mouse models of lymphoma, breast cancer, and melanoma. We describe our protocol for the conjugation of antibodies that are optimized for IF staining of murine tissues preserved as FFPE and provide our protocols for PhenoCycler staining and open-source data analysis, which enables visualization of staining, cell segmentation, cell classification, and neighbourhood/proximity analysis. The protocols described below are tunable and offer flexibility to researchers who wish to use their own antibodies of interest for highly multiplexed staining.

## Results

### Development of a Tunable PhenoCycler Antibody Panel for Staining Murine FFPE Tissue

Our tunable PhenoCycler workflow has four major components: 1) antibody selection; 2) antibody conjugation and optimization; 3) tissue staining and imaging; and 4) data analysis (**Figure 1**).

**Figure 1:**
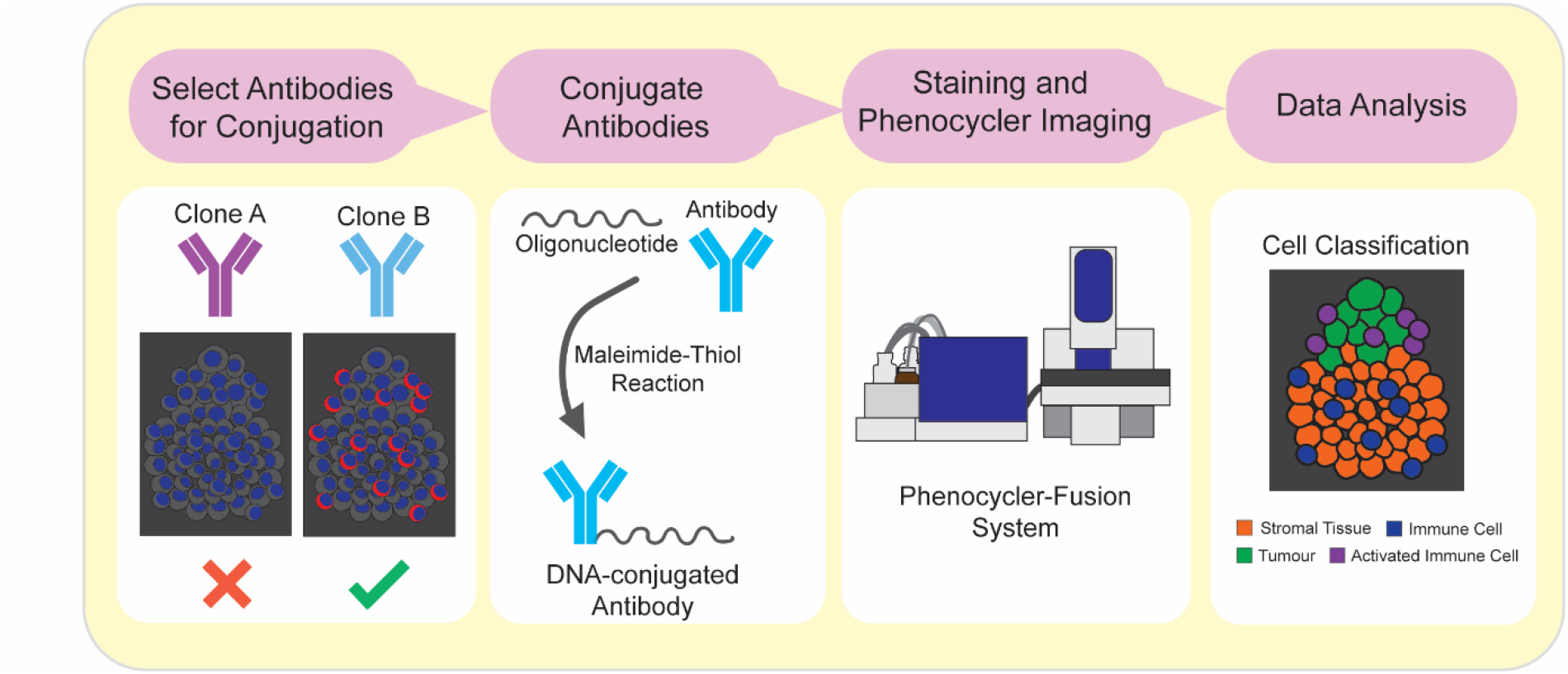
Workflow for selection of antibodies, antibody conjugation, and PhenoCycler staining. Schematic showing the workflow outlined in this Research Resource.

Using the protocols described below, we selected 16 antibodies which could be used to phenotype most common cells found in the murine TME (**Figure 2A**). Each of these antibodies were conjugated to Akoya PhenoCycler barcodes (**Table 1**) and were optimized for PhenoCycler staining. Each barcode has a complementary reporter conjugated to either ATTO550, AF647, or AF750, and barcodes were selected for each antibody with this in mind. In general, antibodies that showed very strong signal-to-noise ratio (SNR) were conjugated to barcodes with AF750-tagged complementary reporters, whereas antibodies that corresponded to antigens of lower abundance and lower expression were conjugated to barcodes with AF647-tagged complementary reporters, and antibodies that marked antigens of medium abundance and weak to medium SNR were conjugated to barcodes with ATTO550-tagged complementary reporters. With this staining panel, we were able to quantify tumour cells, endothelial cells, fibroblasts, myeloid cells (macrophages, neutrophils, and dendritic cells) and lymphoid cells (helper T cells, cytotoxic T cells, regulatory T cells, and B cells) in the murine TME (**Figure 2B**). Furthermore, the protocols for analysis described below can be used to examine how the spatial relationships between these cell types change across tumour models and experimental conditions.

**Figure 2:**
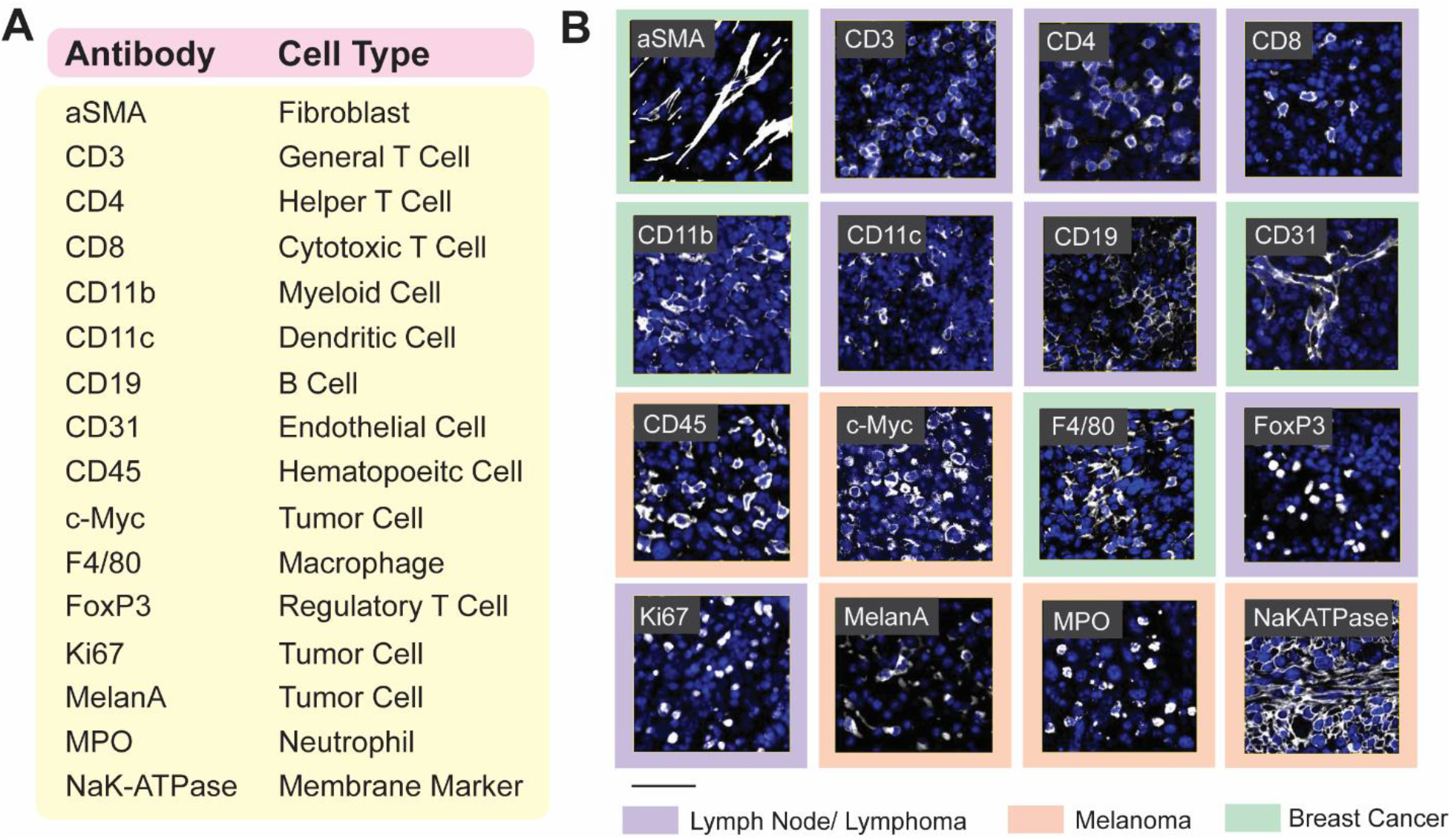
16-plex PhenoCycler staining of murine FFPE tissues. **A.** Table showing the antibodies included in our Murine FFPE PhenoCycler staining panel, and the cell type they are used to identify. **B.** Images showing successful PhenoCycler staining of each antibody in the staining panel. In each image, DAPI is blue, and each individual marker is white. The colour of the outer border indicates the tissue type in the image. Scale bar is 50 µM.

**Table 1.** Primary Antibody Table.

| **Antibody** | **Company** | **Barcode** | **Reporter** |
| --- | --- | --- | --- |
| ***Primary Antibodies*** |  |  |  |
| **Alpha smooth muscle actin (αSMA)** | Abcam | BX014 | RX014-ATTO550 |
| **CD3** | Abcam | BX017 | RX017-ATT0550 |
| **CD4** | Invitrogen | BX002 | RX002-ATTO550 |
| **CD8** | Invitrogen | BX005 | RX005-ATTO550 |
| **CD11b** | Abcam | BX003 | RX003-AF647 |
| **CD11c** | Cell Signaling | BX015 | RX015-AF647 |
| **CD19** | Cell Signaling | BX027 | RX027-AF647 |
| **CD31** | Dianova | BX026 | RX026-ATT0550 |
| **CD45** | R&D Systems | BX007 | RX007-AF750 |
| **c-Myc** | Abcam | BX001 | RX001-AF750 |
| **F4/80** | Cell Signaling | BX020 | RX020-ATTO550 |
| **FoxP3** | Cell Signaling | BX019 | RX019-AF750 |
| **Ki67** | Akoya | BX047 | RX047-ATTO550 |
| **MelanA** | Abcam | BX004 | RX004-AF750 |
| **MPO** | R&D Systems | BX013 | RX013-AF750 |
| **NaK-ATPase** | Abcam | BX023 | RX023-ATTO550 |

### Generation of a Multi-Cancer TMA for PhenoCycler Staining

Given that there is conservation amongst the cell types found in the TME across a number of tumour types (38), we generated a TMA with tumour cores banked from widely used pre-clinical mouse models of lymphoma, breast cancer, and melanoma, and matched normal tissues, with the goal of performing spatial phenotyping of the murine TME. To achieve this, archival FFPE tissue blocks were sectioned and stained with H&E and an anti-CD45 antibody to facilitate selection of immune-rich regions within the tumours for core-punching (**Figure 3A**). From each tissue block, two to three 1 mm cores were included, for a total of 84 cores (**Figure 3B**).

**Figure 3:**
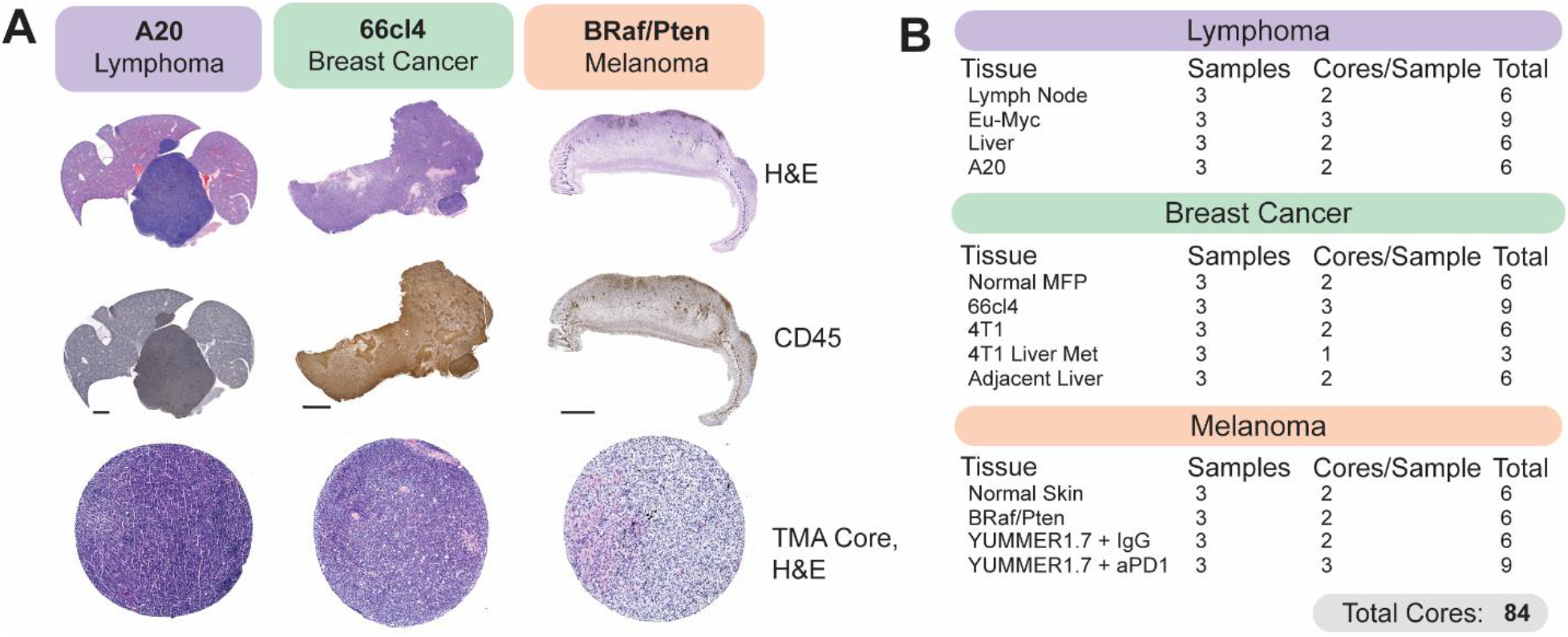
Generation of a murine tissue microarray (TMA) for PhenoCycler Staining. **A.** Representative H&E and CD45 staining from murine tumour tissues. H&E and CD45 staining was used to select regions of interest for TMA core punching. Scale bars represent 100 µm. Bottom row shows H&E staining of the tissue cores, following TMA generation. Each TMA core is 1mm in size. **B.** Table indicating the types and numbers of cores included in our multi-cancer murine TMA.

For the lymphoma portion of the TMA, cores from A20 and Eµ-Myc tumours were included. A20 is a commonly used mouse model of B Cell Non-Hodgkin’s Lymphoma (B-NHL), syngeneic to BALB/C mice (39). Upon tail vein injection, A20 cells will home to the liver to form an aggressive extranodal lymphoma, and samples from day-27 post-A20 tail vein injection were included in the TMA, with matched adjacent non-tumour bearing liver tissue (ie, tissue from a non-tumour bearing liver lobe). Eµ-Myc is a B-NHL model syngeneic to C57BL/6J mice, which forms tumours primarily in the spleen and cervical and inguinal lymph nodes. Samples from the lymph nodes of non-tumour bearing mice and from the cervical lymph nodes of mice at day-14 post-Eµ-Myc injection were included in the TMA.

Tumour samples grown from the 66cl4 and 4T1 murine triple-negative breast cancer cell lines were included in the multi-cancer TMA. Both cell lines are capable of forming primary tumours following inoculation into the mammary fat pads of syngeneic BALB/c mice (40). However, they differ in their metastatic potential and route of dissemination (41). 66cl4 cells are weakly metastatic and tend to travel via the lymphatic system to the lung (41). Samples from our previously published (42) cohort of 66cl4 tumours from day-33 post-injection (roughly 1750 mm^3^ in size) were included. The highly aggressive 4T1 model is metastatic to the bone, lung and liver and predominantly does so via the vasculature (41) (43). We included samples from primary 4T1 tumours harvested day-10 post-injection, when they are 600 mm^3^. Additionally, to define differences between the TME of primary and metastatic 4T1 tumours, samples were included from mice with 4T1 liver metastases, generated using the intrasplenic injection model of experimental metastasis (44).

Finally, to enable comparison of ICI-resistant and ICI-susceptible murine melanoma models, melanomas from the *Tyr::CreER*/*BRaf^CA/+^/Pten^lox/lox^* conditional melanoma model (45) and the YUMMER1.7 syngeneic melanoma model (46) were included. The *Tyr::CreER*/*BRaf^CA/+^/Pten^lox/lox^* transgenic mouse is a well-described murine model of melanoma, which allows 4-hydroxytamoxifen-inducible melanocyte-targeted *BRAF^V600E^* expression and simultaneous *PTEN* inactivation (referred to hereafter as *BRAF^V600E^*/*PTEN^-/-^*). Murine *BRAF^V600E^*/*PTEN^-/-^* melanomas are characterized by low immune cell infiltration and are therefore known to be “immune cold” and resistant to ICI-therapy (47, 48). YUMMER1.7 cells were derived from a *BRAF^V600E^*/*PTEN^-/-^* melanoma following subsequent exposure to ultraviolet radiation to increase mutational burden, making YUMMER1.7 melanomas sensitive to ICI treatment (46). We included samples harvested at 2000 mm^3^ from YUMMER1.7 melanomas treated with either αPD-1 immunotherapy or IgG control.

### Comparing the TME of Nodal and Extranodal Murine B Cell Non-Hodgkin’s Lymphoma

B-NHL is the most commonly diagnosed lymphoid malignancy, arising from the abnormal proliferation of B lymphocytes. B-NHL frequently arises in secondary lymphoid organs, such as the lymph nodes or spleen, but extranodal involvement is common and has been shown to correlate with adverse outcomes (49). Studies have demonstrated that B-NHL has distinct biological features between different extranodal sites (50–52), and mouse modelling provides the opportunity to functionally examine how varied TMEs can impact the B-NHL immune cell infiltration, specifically as A20 tumours develop in the murine liver (**Figure 4A**), while Eµ-Myc tumours develop in the lymph nodes (**Figure 4B**).

**Figure 4:**
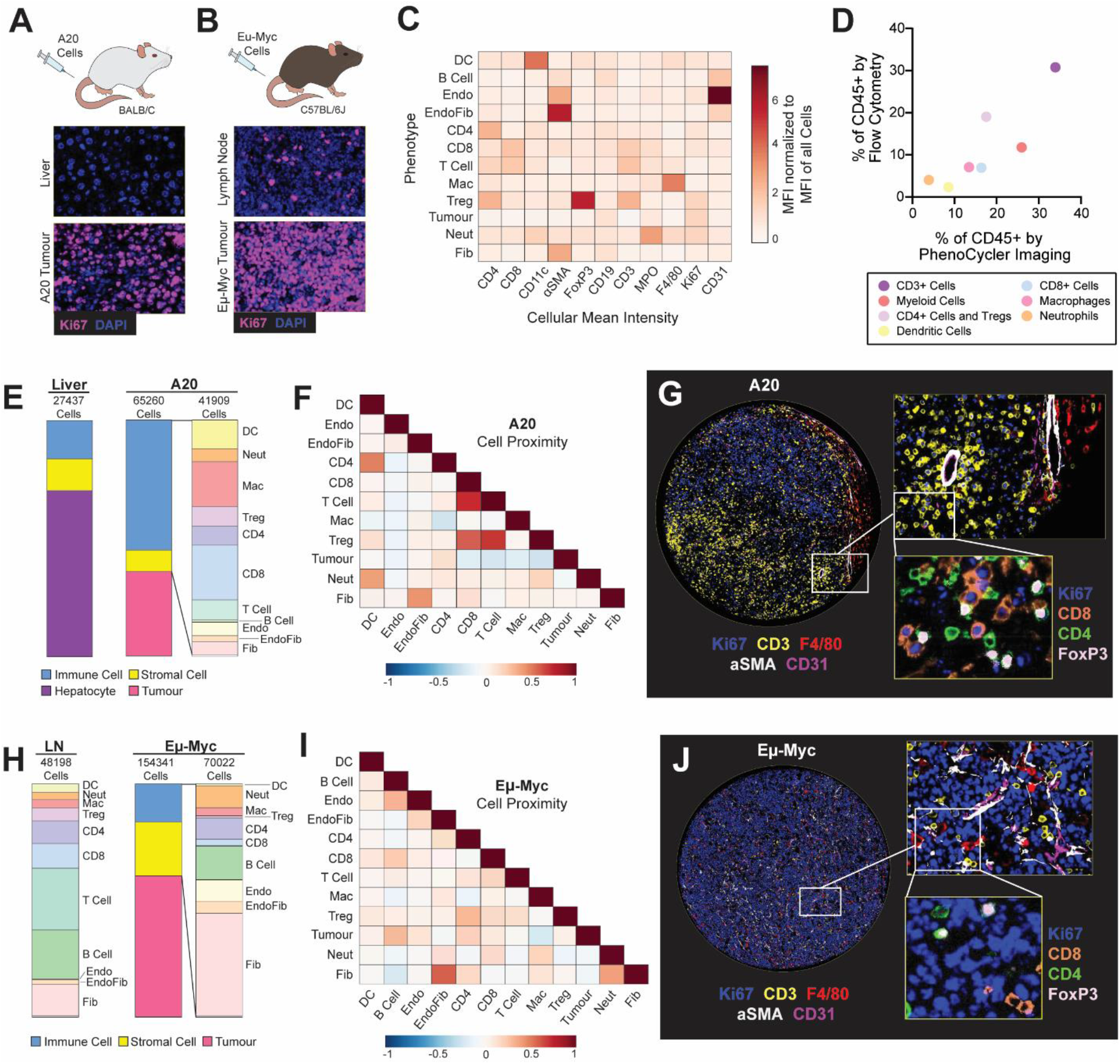
PhenoCycler imaging of the murine nodal and extra-nodal B-cell lymphoma tumour microenvironment. **A.** Schematic of the A20 model of extranodal B-NHL, and representative images of Ki67 staining in healthy liver and A20 tumour-bearing liver. **B.** Schematic of the Eµ-Myc model of nodal B-NHL, and representative images of Ki67 staining in healthy lymph node and an Eµ-Myc tumour-bearing lymph node. **C.** Heatmap showing the normalized cellular mean intensity of markers within the PhenoCycler staining panel, in different phenotypes of cells in A20 and Eµ-Myc tumours. **D.** Scatter plot comparing the proportions of different cell phenotypes as determined by PhenoCycler staining versus archival flow cytometry data, for A20 tumours. Pearson r = 0.8551, p = 0.0142. **E**. Proportions of different cell types in adjacent healthy liver and A20 tumour-bearing liver. **F.** Heatmap showing neighborhood analysis of A20 tumours, as Pearson correlation coefficient between cells. Blue hue indicates cells are likely to be in further proximity, while red hue indicates that cells are likely to be in closer proximity. **G.** Representative image of an A20 tumour core. **H.** Proportions of different cell types in healthy lymph nodes and Eµ-Myc tumour-bearing lymph nodes. **I.** Heatmap showing neighborhood analysis of A20 tumours, as Pearson correlation coefficient between cells. **J.** Representative image of an Eµ-Myc tumour core.

Following PhenoCycler staining, DAPI-based segmentation of images was performed to extract single-cell marker expression, and cells were classified into phenotypes based on marker expression (**Figure 4C**; see protocols below). In A20 and Eµ-Myc tumours, we were able to detect dendritic cells, B cells, endothelial cells, CD4+ T cells, CD8+ T cells, macrophages, regulatory T cells (Tregs), tumour cells, neutrophils, and fibroblasts. Of note, in these tissues and in the tissues derived from other tumour types, CD31+ endothelial cells formed close contacts with αSMA+ fibroblasts, leading to fluorescence spillover of CD31 and αSMA lineage markers following cell segmentation. We classified these cells as “EndoFib”, representing close contacts between endothelial cells and fibroblasts. This was similarly observed with tightly packed CD4+ and CD8+ T cells in lymphoma tissues only, and we termed these cells “T Cells” in downstream analyses. Despite these challenges in cell segmentation, the proportions of immune cell types found in the A20 TME by PhenoCycler correlated closely with archival flow cytometry immunophenotyping of dissociated A20 tumours, showing that these two methodologies can similarly identify cells in the TME (**Figure 4D**).

The A20 B-NHL TME was characterized by high infiltration of immune cells (55.12%), relative to adjacent non-tumour bearing liver (16.27%) (**Figure 4E**). The A20 immune infiltration was comprised of dendritic cells, macrophages, CD8+ T cells, and Tregs, while immune cells in the adjacent liver were almost exclusively macrophages (likely Kupffer cells), consistent with what is expected in normal liver. We analyzed spatial interactions between the different cell phenotypes in A20 tumours using CytoMAP to calculate the probability of different cell types being within 50 µM of each other (53) (see methods). We found that Tregs were in close proximity to T cells (correlation coefficient = 0.695) and CD8+ T cells (correlation coefficient = 0.6011). Furthermore, tumour cells were spatially segregated from immune cells such as CD8 T cells (correlation coefficient = -0.1289), macrophages (correlation coefficient = -0.1287), and Tregs (correlation coefficient = -0.1942; **Figure 4F-G**). These results suggest that tumour cells tend to localize together within the extranodal B-NHL tumour mass while immune cells localize together at the tumour periphery and highlight that Tregs are a critical mediator of CD8+ T cell immunosuppression in A20 tumours.

As expected, non-tumour bearing murine cervical lymph nodes consisted almost entirely of immune cells (84.04%); however, the presence of Eµ-Myc tumours drastically decreased this proportion (16.41%). In Eµ-Myc tumours, the overall immune composition was altered relative to healthy lymph nodes, with an increase in neutrophils, and a decrease in Tregs, dendritic cells, and CD8+ T cells (**Figure 4H**). Eµ-Myc tumours also had an increased proportion of stromal cells, including fibroblasts and endothelial cells (23.09% in Eµ-Myc tumours, compared to 15.96% in healthy lymph node). Spatial analysis further demonstrated that Eµ-Myc tumours are relatively disorganized (**Figure 4I**), and different cell types seem to be randomly distributed throughout the tumour. For instance, while CD8+ T cells and Tregs can be detected (**Figure 4J**), they are spatially segregated and are likely not functionally interacting (correlation coefficient = 0.2058).

Our data shows that the presence of A20 extranodal tumours induces the recruitment of immune cells to the liver, while the presence of Eµ-Myc nodal tumours leads to immune cell displacement from the lymph nodes. Furthermore, as it has been previously suggested (54), our data suggest that A20 tumours rely on Tregs to induce immunosuppression and achieve immune evasion, while Eµ-Myc tumours are immune-depleted, and therefore do not require inhibitory immune cell interactions to achieve immunosuppression. Thus, these two models of B-NHL employ drastically different strategies to avoid immune destruction.

### Defining Differences in the TME of Primary and Metastatic Murine Breast Cancer

Breast cancer is a heterogeneous disease, comprised of different molecular subtypes. Patients with triple-negative breast cancer (TNBC) have the worst prognosis, largely due to aggressive tumour behaviour, increased risk of metastasis, and resistance to conventional anti - cancer therapies (55). Treatments which target the TME in TNBC have gained increased attention in recent years, spurred on by data demonstrating the strong immunogenicity of this tumour type (56) and success of combined chemotherapy and immunotherapy in clinical trials (57, 58). Understanding the cellular landscape of TNBC tumours will undoubtedly be beneficial for the continued development of successful TME-targeting therapies.

Towards this goal, we used PhenoCycler to image primary tumours from the commonly used pre-clinical murine 66cl4 and 4T1 TNBC models, as well as 4T1 liver metastases (**Figure 5A-B**). Using the protocols described below, we performed cell-segmentation and cell-clustering to identify cell phenotypes. In these tumours, we could identify the same immune and stromal cell types as were found in lymphoma tumours. However, while lymphoma tumour cells were characterized by Ki67 positivity, we found that tumour cells in breast cancer models could be stratified based on Ki67 expression (**Figure 5D**), and both Ki67+ and Ki67-tumour cells were numerous enough to merit individual classification. Interestingly, the percentage of Ki67+ tumour cells was higher in the more aggressive 4T1 samples compared to 66cl4 (**Figure 5E**; percentage of Ki67+ tumour cells among total tumour cells: 66cl4: 41.13%; 4T1: 80.65%; 4T1-liver: 63.47%). The proportion of CD45+ immune cells was similar in all tumour sample types (**Figure 5E**; 66cl4: 45.39%; 4T1-primary: 52.52%; 4T1-liver: 49.84%), with macrophages representing the dominant immune cell type (**Figure 5E-F**: 66cl4: 39.35%; 4T1-primary: 40%; 4T1-liver: 38.59%) in line with previously published reports (59).

**Figure 5:**
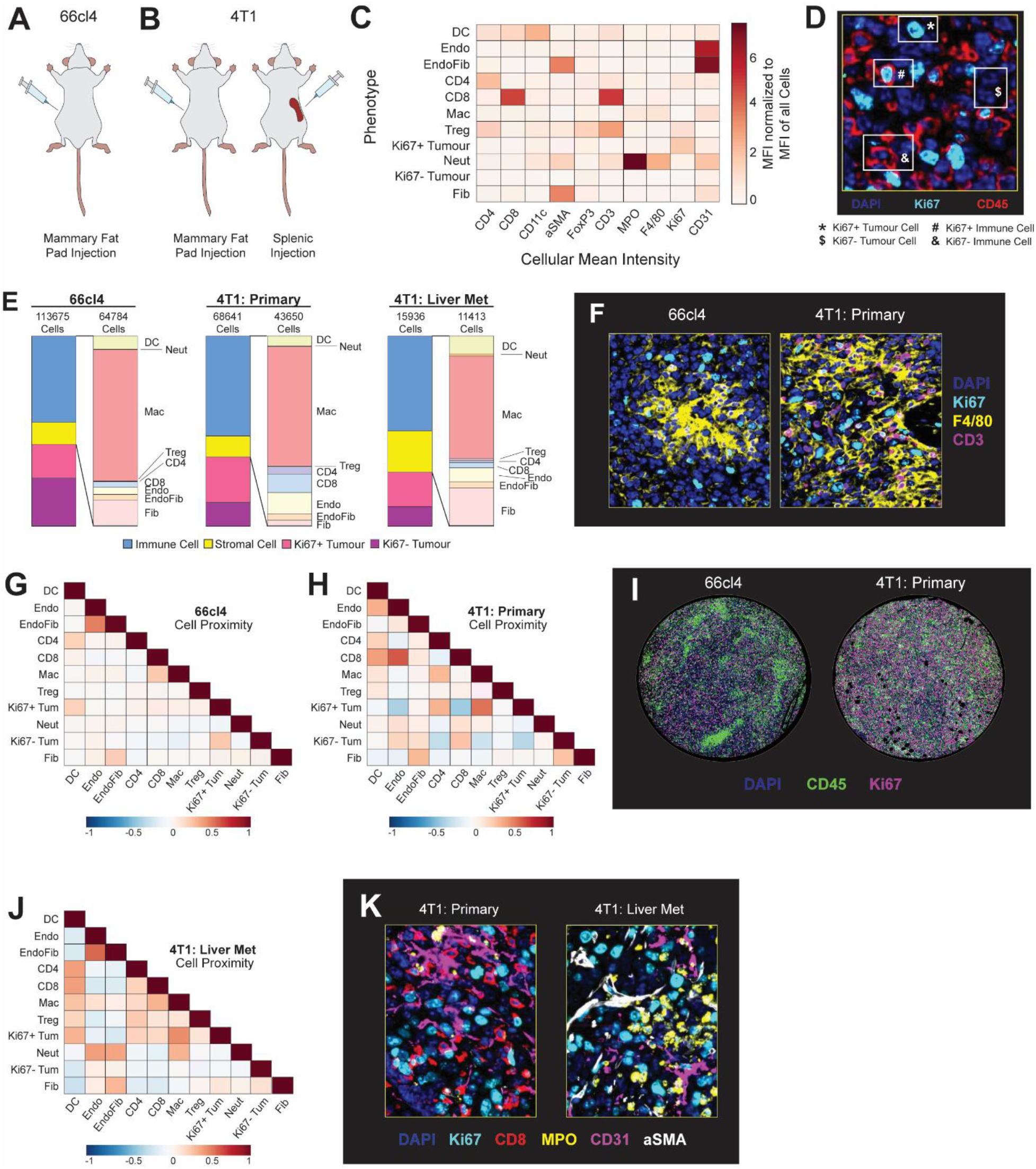
PhenoCycler imaging of the murine breast cancer tumour microenvironment. **A.** Schematic of the 66cl4 murine model of breast cancer. **B.** Schematics of the 4T1 murine models of breast cancer and breast cancer liver metastasis. **C.** Heatmap showing the normalized cellular mean intensity of markers within the PhenoCycler staining panel, in different phenotypes of cells in 66cl4 and 4T1 tumours**. D.** Representative image showing Ki67+ and Ki67-tumour cells. **E.** Proportions of different cell types in 66cl4 and 4T1 tumours. **F.** Representative images of macrophages and T cells in 66cl4 and 4T1 primary tumours.. **G.** Heatmap showing neighborhood analysis of 66cl4 tumours. as Pearson correlation coefficient between cells. **H.** Heatmap showing neighborhood analysis of 4T1 primary tumours, as Pearson correlation coefficient between cells. **I.** Representative images showing immune cell infiltration patterns in 66cl4 and 4T1 tumours. **J.** Heatmap showing neighborhood analysis of 4T1 liver metastases, as Pearson correlation coefficient between cells. **K.** Representative images comparing immune and stromal cell types in 4T1 primary and 4T1 liver metastases.

In addition to the composition of the immune cell landscape, cell neighbourhood analyses highlighted further differences between tumour types. Immune cells in 66cl4 tumours were largely localized together in restricted regions, but were found to be intermingling with other cell types throughout 4T1 tumours (**Figure 5G-I**). In particular, 4T1 tumours were observed to have strong spatial interactions between CD8+ T cells and endothelial cells (correlation coefficient = 0.6198), and Ki67^+^ tumour cells and macrophages (correlation coefficient = 0.5448; **Figure 5H**). In contrast, the interaction between endothelial cells and CD8+ T cells is lost in 4T1 liver metastases (correlation coefficient = -0.1667) compared to the primary tumour, with a concomitant increase in interactions between endothelial cells and neutrophils (correlation coefficient = 0.4064) and total neutrophil abundance (**Figure 5J-K**; 4T1: 0.19%; 4T1-liver: 0.88%). These data corroborate observations that formation of 4T1 liver metastases is heavily reliant on the infiltration of neutrophils into the TME (60), suggesting that proximity to the vascular endothelium may be indicative of immune cell influx patterns.

These data illustrate the utility of PhenoCycler technology to profile the immune landscape of murine TNBC tumours, as we characterized immune cell composition of FFPE-processed murine tumour types while layering on top cellular distributions in space. We propose that future applications of PhenoCycler technology, using in-depth antibody panels which assess immune cell function or polarization, may aid in uncovering therapeutic options to augment anti-tumour immunity in TNBC patients.

### Characterizing the TME of ICI-Resistant and ICI-Susceptible Murine Melanoma

Melanoma is one of the most immunogenic cancer types, due to its high mutational burden, which leads to the production of neoantigens that are recognized by patrolling immune cells. To this end, ICI therapies have revolutionized the treatment of melanoma, but innate and acquired resistance remain as clinical challenges. Furthermore, clinical studies have shown that ICI resistance is associated with changes in TME composition (61, 62).

We used two immune competent murine models of melanoma for PhenoCycler staining: the *BRAF^V600E^*/*PTEN^-/-^*model and the YUMMER1.7 model (**Figure 6A**). *BRAF^V600E^*/*PTEN^-/-^*melanomas exhibit high intratumoural heterogeneity and melanoma cell plasticity, are known to be immune “cold”, and are insensitive to ICI treatment. Conversely, YUMMER1.7-derived tumours are highly immunogenic and are susceptible to ICI-mediated tumour inhibition (46). In our previous work, we have shown that αPD-1 immunotherapy reduced the growth of YUMMER1.7 tumours and improved the overall survival of mice, but most tumours failed to go into complete remission (48), mimicking the human clinical scenario where more than half of patients experience disease progression following αPD-1 treatment (63). Thus, we aimed to determine if tumour regrowth following αPD-1 treatment is associated with TME remodeling by comparing isotype control (IgG)-treated tumours with αPD-1-treated tumours (αPD-1-relapsed), harvested when tumours were 2000 mm^3^. Additionally, samples from *BRAF^V600E^*/*PTEN^-/-^* tumours facilitated further comparison between an ICI-resistant and an ICI-sensitive murine model of melanoma.

**Figure 6:**
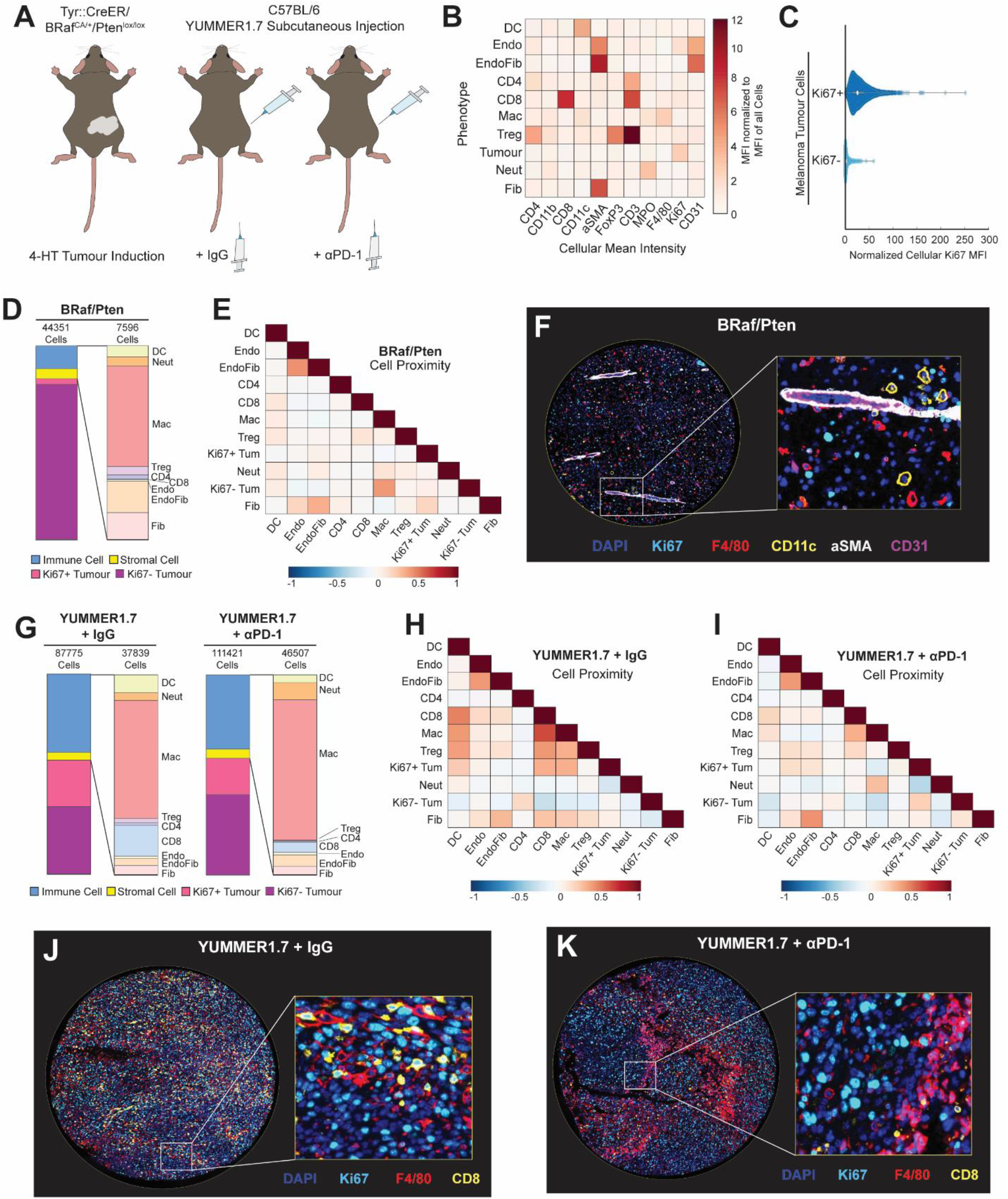
PhenoCycler imaging of the murine melanoma tumour microenvironment. **A.** Schematics of the BRaf/Pten and YUMMER1.7 murine models of melanoma. **B.** Heatmap showing the normalized cellular mean intensity of markers within the PhenoCycler staining panel, in different phenotypes of cells in BRaf/Pten and YUMMER1.7 tumours. **C.** Normalized Ki67 mean fluorescence intensity in Ki67+ tumour cells and Ki67-tumour cells. **D.** Proportions of different cell types in BRaf/Pten tumours. **E.** Heatmap showing neighborhood analysis of BRaf/Pten tumours, as Pearson correlation coefficient between cells. **F.** Representative image of BRaf/Pten tumour core. **G.** Proportions of different cell types in YUMMER1.7 tumours, treated with IgG control or αPD-1. **H-I.** Heatmap showing neighborhood analysis of YUMMER1.7 tumours treated with IgG control (**H**) or αPD-1 (**I**), as Pearson correlation coefficient between cells. **J.** Representative image of YUMMER1.7-IgG tumour core. **K.** Representative image of YUMMER1.7-αPD-1 tumour core.

PhenoCycler images from these murine melanomas were cell-segmented and classified based on marker expression (**Figure 6B**). Similarly to breast cancer, we found that two distinct populations of tumour cells were present: Ki67+ and Ki67-(**Figure 6C**). *BRAF^V600E^*/*PTEN^-/-^*tumours were composed of 79.98% Ki67-tumour cells, and 2.88% Ki67+ proliferating tumour cells (**Figure 6D**). These data are consistent with our previous work demonstrating that *BRAF^V600E^*/*PTEN^-/-^*melanoma cells typically undergo phenotype switching from a more proliferative to a more invasive state, that is characterized by slower proliferation. The remaining 17.12% of cells within *BRAF^V600E^*/*PTEN^-/-^*tumours were stromal cells (5.29%) and immune cells (11.83%). The majority of the immune cells were found to be macrophages, with minimal T cell infiltration. Consistent with the fact that *BRAF^V600E^*/*PTEN^-/-^*tumours are immune “cold”, spatial analysis demonstrated that cells within these tumours did not have preferential interactions with each other (**Figure 6E**) and appeared randomly distributed within the tissues (**Figure 6F**).

As compared to *BRAF^V600E^*/*PTEN^-/-^* tumours, YUMMER1.7 tumours had a significantly higher proportion of proliferative (Ki67+) tumour cells, which was slightly decreased upon resistance to αPD-1 treatment (**Figure 6G**; 23.23% in IgG-treated samples versus 18.22% in αPD-1-relapsed samples). Furthermore, both IgG-treated and αPD-1-relapsed YUMMER1.7 tumours were more immunogenically “hot” with increased immune cell abundance as compared to *BRAF^V600E^*/*PTEN^-/-^* tumours (IgG-treated: 39.11% immune cells; αPD-1-relapsed: 37.1% immune cells). Whie IgG-treated and αPD-1 relapsed YUMMER1.7 tumours had similar immune cell invasion, there were distinct differences in cellular organization. Spatial analysis of YUMMER1.7-IgG tumours showed strong interactions between CD8+ T cells and macrophages (correlation coefficient = 0.6482), and CD8+ T cells and dendritic cells (correlation coefficient = 0.4957; **Figure 6H**). YUMMER1.7-IgG Ki67+ tumour cells were in close proximity to these immune cells (CD8+ T cell/ Ki67+ tumour cell correlation coefficient = 0.3529; macrophage/ Ki67+ tumour cell correlation coefficient = 0.3535; dendritic cell/ Ki67+ tumour cell correlation coefficient = 0.2486), as compared to Ki67-tumour cells (CD8+ T cell/ Ki67-tumour cell correlation coefficient = -0.2316; macrophage/ Ki67-tumour cell correlation coefficient = -0.1342; dendritic cell/ Ki67-tumour cell correlation coefficient = -0.1013). However, Ki67-tumour cells were in closer contact with CD4+ T cells (Ki67+ tumour cell/ CD4+ T cell correlation coefficient = -0.0339, Ki67-tumour cell/ CD4+ T cell correlation coefficient = 0.1932). In αPD-1-relapsed tumours, all of these cellular contacts were reduced (**Figure 6I-K**), supporting reduced tumour-immune cell interaction as a mechanism of acquired ICI resistance in melanoma.

All together, these results support that Ki67+ proliferative melanoma cells have higher immunogenicity. In agreement with this, *BRAF^V600E^*/*PTEN^-/-^*tumours have a substantially increased proportion of Ki67-tumour cells, correlating with a decreased proportion of infiltrating immune cells. Moreover, in YUMMER1.7-IgG tumours, Ki67+ tumour cells maintain close contacts with immune cells. In αPD-1-replapsed YUMMER1.7 tumours, there is no preferential interaction of Ki67+ or Ki67-tumour cells with immune cells, indicating immune dysfunction upon the emergence of ICI-resistance. To this end, our data supports the notion that ICI-resistance is associated with decreased interactions between immune cells and tumour cells (64), as αPD-1-relapsed YUMMER1.7 tumours have similar macrophage infiltration as compared to IgG controls, yet the tissue organization is altered such that there are limited cellular contacts between macrophages and tumour cells.

### Discussion

#### PhenoCycler Imaging of Murine FFPE Tumour Tissues

The TME is a central player in many of the biological challenges associated with cancer treatment, such as immune escape, disease metastasis, and drug resistance. Thus, it is critically important to assess both the composition and the spatial dynamics of the TME in mouse models that are commonly used in pre-clinical cancer research. Previously, PhenoCycler imaging of murine tissues had been limited to fresh frozen tissues. Here, we detail imaging FFPE murine tissues and provide our protocols for the optimization and conjugation of antibodies for this purpose. To illustrate the feasibility of this approach, we provide data showing successful staining of murine lymphoma, melanoma, and breast cancer tissues.

Immunofluorescence imaging of FFPE tissues is not without challenges. FFPE tissues tend to have high auto-fluorescence, which can distort true positive staining. Additionally, formalin-fixation induces protein cross-linking, leading to epitope masking and difficulties in primary antibody binding (65). However, many research groups archive tissues from previous pre-clinical studies in FFPE format; thus, it is a worthwhile endeavor to optimize antibodies for highly multiplexed imaging of murine FFPE tissues, to allow for the utilization of archival materials. To this end, the selection of antibody clones with an ideal SNR was a critical first step towards this goal. Following clone selection, antibodies were carefully optimized, for parameters such as concentration, incubation time and temperature, and imaging exposure time.

In this dataset, we first showed that the tumour microenvironment of murine B-NHL is altered between the A20 and Eµ-Myc models of B-NHL, suggesting two different mechanisms of immune evasion. Then, we demonstrated that the distribution of the immune microenvironment differs between models of murine TNBC, and showed how measurement of interactions between endothelial and immune cells may relate to TME infiltration. Finally, using samples from murine melanoma, we examined how the TME is altered in the context of ICI-resistance, and found that ICI-susceptible tumours have increased spatial interactions between immune cells and tumour cells. Our data asserts that the careful selection of a mouse model is critical when designing experiments to study the TME. For instance, Eu-Myc or *BRAF^V600E^*/*PTEN^-/-^*models may be appropriate to study therapeutics that are predicted to increase immune cell trafficking or retention in the TME; while A20, 66cl4, 4T1, or YUMMER1.7 models could be useful to study therapeutics that re-activate immune cells already present in the TME. Furthermore, we demonstrate that PhenoCycler imaging of murine tumours can be employed both to test and to generate hypotheses. As an example of this, we hypothesized that the TME would be altered in different models of B-NHL, and our data found close cellular contacts between CD8+ T cells and Tregs in A20 B-NHL tumours, but not in Eµ-Myc tumours. Thus, one may hypothesize that Tregs in A20 function via direct inhibitory interactions with CD8+ T cells to suppress anti-tumour immunity (66), and to further investigate this, *ex vivo* functional assays could be employed. Throughout this study, there are numerous examples where our findings via PhenoCycler imaging have been hypothesis generating and could be further explored with i*n vitro* or *in vivo* experimentation.

#### Analysis of Highly Multiplexed Immunofluorescence Staining Data

While many labs may be eager to begin highly multiplexed imaging of their experimental tissues, data analysis can appear to be a daunting task. Below, we provide our workflows for open-source analysis of PhenoCycler imaging data. In our analysis pipeline, we primarily use QuPath software for cell classification (67), and CytoMAP for spatial analysis (53). In QuPath, images are segmented into single cells using a StarDist plugin (68, 69). In some cases, cell segmentation failed to discriminate individual cells when close contacts resulted in fluorescence spillover. This was particularly true in the case of intact blood vessels, where αSMA+ fibroblasts formed close contacts with CD31+ endothelial cells. In our dataset, we referred to these as “EndoFib” cells, and considered them to be a distinct entity. We also note that alternate segmentation methods that incorporate a cell membrane marker to define cellular boundaries may need to be utilized when the primary cell type of study is irregularly shaped or multinucleated, such as a fibroblast or a neuron (70).

To classify cells into phenotypes, we manually annotated a small number of cells based on their marker expression and used object-based classification methods in QuPath to extend this cell classification to the whole tissue. While this method of analysis proved to be highly successful in our hands, other analysis pipelines may allow more cursory or in-depth higher-plex image analysis. For instance, following cell segmentation, cellular mean intensity of all markers can be exported to a comma-separated values (CSV) file, which can be analyzed with FlowJo or other programs (so-called “hand-gating”). However, the success of hand-gating is limited by cell segmentation noise (71). Another alternative is to perform unsupervised clustering analysis, using pipelines such as Seurat, but we note that over-clustering has the potential to identify false phenotypes, and therefore must be used with caution. Overall, the analysis pipeline described below is an excellent starting point for novices in multiplexed immunofluorescence image analysis and can be built upon to allow for more sophisticated analyses which answer increasingly complex experimental questions.

#### Limitations of the Technology

While the PhenoCycler system for highly multiplexed fluorescent imaging has distinct advantages over other highly multiplexed imaging platforms, such as non-destructive tissue imaging, limited spectral overlap in fluorescence due to iterative cycles of imaging, and the use of robotic automation to increase throughput, there are also limitations to this technology. For instance, it is expensive and time consuming to identify antibody clones that are suitable for PhenoCycler immunofluorescence imaging. Additionally, the conjugation of an antibody to a DNA barcode can occasionally result in antibody dysfunction, and it is costly to research labs to correct problems of this nature. The process of identifying antibody clones suitable for PhenoCycler imaging represents a significant bottleneck in the PhenoCycler workflow, especially when generating custom antibody panels.

Furthermore, while PhenoCycler has been proven to image up to 100 markers, there is limited opportunity for signal amplification to aid in the visualization of targets of low abundance. To this point, there have been attempts to integrate tyramide-based signal amplification into the PhenoCycler workflow (20), but the proposed strategy requires iterative staining and stripping cycles, thereby increasing the risk of tissue damage and decreasing automation.

#### Concluding Remarks

As new technologies in highly multiplexed imaging continue to emerge, we predict that many labs will require refined protocols for image acquisition and data analysis. Highly multiplexed imaging provides the opportunity to visualize many diverse cell types in their native environments, and the insights provided from these types of experiments are instrumental in advancing the field of cancer research. Thus, we predict that the number of publications which employ highly multiplexed imaging will explode over the next decade. To this end, data must be appropriately collected and analyzed, and we hope to empower research groups to begin working towards this goal with the protocols provided below.

## Materials and Methods

### Selection and Validation of Antibodies for Conjugation, and Quality Control of Staining

Due to epitope masking associated with FFFPE preservation (65), the careful selection of antibodies is critical to successful PhenoCycler staining. Below, we describe our IF staining protocol for the selection of antibody clones which can prioritized for barcode conjugation. All antibodies should be tested on the tissue they are ultimately meant to stain.

#### Deparaffinization and Antigen Retrieval

1. Mount 4 µm microtome tissue sections onto SuperFrost Plus slides (Fisherbrand).
2. Deparaffinize slides using the following solutions, for 5 minutes each: Xylene (1), Xylene (2), 100% EtOH (1), 100% EtOH (2), 95% EtOH, 70% EtOH, 50% EtOH, and running tap water.
3. Transfer slides to a PT Link Pre-treatment machine filled with <u>1X Tris-EDTA antigen</u> <u>retrieval buffer (pH 9.0) and</u> cook at 90°C for 20 minutes. After depressurization, cool slides for 1 hour.

i. Note: Less toxic alternatives, such as HistoChoice, can be used in place of Xylene.
ii. Note: Recipes for all solutions used in these protocols can be found in **Table 2**. Recipes listed below will be <u>underlined</u>.

#### Blocking

4. Rinse slides in tap water and dry the glass around the tissue with a Kimwipe. Circle tissue with a hydrophobic PAP pen, and rinse with 2 changes of <u>IF Wash Buffer</u>.
5. Block slides for 30 minutes at room temperature with <u>Primary Blocking Buffer</u>, then rinse with 2 changes of <u>IF Wash Buffer</u>.
6. Block slides for another 30 minutes at room temperature with <u>FC Blocking Buffer</u>, then rinse with 2 changes of <u>IF Wash Buffer</u>.

#### Primary and Secondary Antibody Incubation

7. Dilute primary antibody in <u>Antibody Buffer</u> and incubate slides in primary antibody at 4°C overnight in a humidity chamber.

i. Note: For initial optimization, we try 10ug/ml antibody dilution (approximatively 1 in 100).
ii. Note: Staining specificity can be improved for some antibodies by incubating with a higher antibody concentration (eg. 20ug/ml), for 30 minutes at 37°C.
8. Rinse slides with 3 changes of <u>IF Wash Buffer</u>.
9. Incubate slides for 1 hour at room temperature with secondary antibody conjugated to AlexaFluor647, diluted 1 in 500 in <u>Antibody Buffer</u>.
10. Rinse slide with 3 changes of <u>IF Wash Buffer</u>.

#### Counterstaining, Mounting, and Imaging

11. Stain tissue with <u>prepared DAPI</u> for 15 minutes, then rinse slide 3 times with <u>IF Wash</u> <u>Buffer</u>.
12. Mount coverslips onto slides with Flouromount-G, and then allow to dry for 15 minutes.
13. Image slides with the same microscope that will be used for PhenoCycler image acquisition.

i. Note: Acquiring on the same microscope used for the Phenocycler image acquisition will give a better representation of the final staining. In this study we used the Fusion microscope from Akoya Biosciences.
ii. Note: The results from optimization staining will help in the subsequent steps in assessing the efficacy of the antibody conjugation by comparing both stains.

#### Assessing IF Staining

Assessing staining quality is challenging. Appropriate negative and positive tissue controls are required. If possible, staining assessment by a pathologist can guide selection of the most appropriate antibody clones. Ideally, a TMA comprising an array of different tissues and pathologies will provide the opportunity for robust assessment of antibody specificity and sensitivity, but whole-tissue slides can be used if a TMA is not available. Critical parameters to consider include:

a. if staining pattern within the tissue consistent with reported literature. Multiple online resources can be used, such as ProteinAtlas.
b. SNR: this parameter will guide the user to which fluorescent reporter to use. For example, if the SNR is very high, the dim AF750 reporter should but used, while the bright AF647 can be used for markers with low SNR.

**Table 2.** Recipes for IF Staining, Antibody Conjugation, and PhenoCycler Staining Solutions.

| **Solution Name** | **Composition** |
| --- | --- |
| **10X Tris-EDTA Antigen Retrieval Buffer, pH 9.0** | 6.05 g Tris<br>1.85 g EDTA<br>400 mL ddH<sub>2</sub>O<br>- Adjust to pH 9.0<br>- Complete to 500 mL with ddH<sub>2</sub>O<br>Store at 4°C for up to <u>30 Days</u> |
| **1X Tris-EDTA Antigen Retrieval Buffer, pH 9.0** | 50 mL 10X Tris/EDTA Buffer pH 9.0<br>450 mL ddH<sub>2</sub>O<br>250 µL Tween20<br>- Mix well and make fresh. |
| **10X Tris Buffered Saline (TBS)** | 80 g NaCl<br>2 g KCl<br>30 g Tris<br>- Adjust pH to 7.4<br>- Complete to 1000 mL with ddH<sub>2</sub>O |
| **IF Wash Buffer** | 200 mL 10X TBS<br>800 mL ddH<sub>2</sub>O<br>250 uL Tween20 |
| **Primary Blocking Buffer** | 1000 µL IF Wash Buffer<br>20 µL Goat <u>or</u> Donkey Serum<br>- Vortex to mix. |
| **FC Blocking Buffer** | 500 µL FC Block<br>5 µL Anti-Mouse HRP<br>- Vortex to mix. |
| **Antibody Buffer** | 1000 µL IF Wash Buffer<br>1 µL Goat <u>or</u> Donkey Serum<br>- Vortex to mix. |
| **Prepared DAPI** | 500 µL PBS<br>1 µL 1mg/mL DAPI<br>- Vortex to mix. |
| **Antibody Reduction Master Mix (A)**<br>**(for 1 conjugation)** | 6.6 µL Reduction Solution 1 (A)<br>275 µL Reduction Solution 2 (A)<br>- Thawed aliquots of Reduction Solution 1 (A) should not be re-used |
| **Bleaching Solution** | 0.8 mL 10M NaOH<br>2.7mL 50% H<sub>2</sub>O<sub>2</sub><br>26.5 mL 1X PBS |
| **Staining Buffer with Blockers (A)**<br>**(for 2 samples)** | 362 µL Staining Buffer<br>9.5 µL N Blocker (A)<br>9.5 µL G2 Blocker (A)<br>9.5 µL J Blocker (A)<br>9.5 µL S Blocker (A) |
| **Post-Staining Fixation (A)** | 1 mL 16% PFA<br>9 mL Storage Buffer (A) |
| **Final Fixative Solution (A)** | 1000 μL 1X PBS<br>20 μL Fixative Reagent (A)<br>- Thawed aliquots of Fixative Reagent (A)<br>should not be re-used |
| **Screening Buffer (A)** | 3.5 mL 10X PhenoCycler Buffer (A)<br>24.5 mL Nuclease-Free Water<br>7 mL DMSO<br>- Allow the Screening Buffer to equilibrate to<br>room temperature prior to use |
| **Reporter Stock Solution (A)**<br>**(for 5 cycles)** | 1220 μL Nuclease Free Water<br>150 μL 10X PhenoCycler Buffer (A)<br>125 μL Assay Reagent (A)<br>5 μL Nuclear Stain (A) |

**Table 3:** Reagents and Tools Table.

| Reagent or Resource | Reference or Source | Identifier or Catalog No. |
| --- | --- | --- |
| ***Experimental Models*** |  |  |
| BALB/C Mice | Charles River | BALB/cAnNCrl |
| C57BL/6J Mice | The Jackson Laboratory | Strain #:000664 |
| Braf/PTEN Mice | The Jackson Laboratory | Strain #:013590 |
| A20 | ATCC | TIB-208 |
| Eμ-Myc | Lab of Dr. Jerry Pelletier | N/A |
| 66cl4 | Lab of Dr. Josie Ursini-Siegel | N/A |
| 4T1 | ATCC | CRL-2539 |
| YUMMER1.7 | Lab of Dr. Marcus Bosenberg | N/A |
| ***Drugs and Treatments*** |  |  |
| 4-Hydroxytamoxifen | Sigma-Aldrich | H6278 |
| IgG Control | Bio X Cell | 2A3, BE0089 |
| aPD-1 | Bio X Cell | RMP1-14, BE0146 |
| ***Chemicals and Reagents*** |  |  |
| Tris | Bio Basic | TB0195 |
| EDTA | Bio Basic | EB0185 |
| Sodium Chloride | Bio Basic | SB0476 |
| Potassium Chloride | Bio Basic | PB0440 |
| Sodium Hydroxide 10N | VWR | BDH7247-1 |
| 50% H<sub>2</sub>O<sub>2</sub> | Sigma-Aldrich | 516813-500ML |
| Paraformaldehyde 16% | Electron Microscopy Sciences | 15710 |
| Tween20 | VWR | 0777-1L |
| ***IHC and IF Reagents and Tools*** |  |  |
| SuperFrost Plus Slides | Fisher | 22-037-246 |
| Xylenes | Fisher | X5-4 |
| Ethanol | Commercial Alcohols | P016EAAN |
| Hydrophobic Barrier PAP Pen | Thermo Scientific | R3777 |
| Harris' Hematoxylin | Sigma-Aldrich | 638A-85 |
| Eosin Y Solution | Sigma-Aldrich | HT110116 |
| Donkey Serum | Jackson ImmunoResearch | 017-000-121 |
| FC Blocking Reagent | Made in house |  |
| ECL Anti-mouse IgG, Horseradish peroxidase linked whole antibody from sheep | Cytiva | NA931V |
| Mouse CD45 Antibody | R&D Systems | AF114 |
| Dnk pAb to Goat IgG (HRP polymer) | Abcam | Ab214881 |
| ImmPACT DAB Substrate Kit, Peroxidase | Vector Laboratories | SK-4105 |
| AF647 donkey anti-rabbit IgG | Invitrogen | A31573 |
| Donkey anti-Rat IgG DyLight 650 | Invitrogen | SA5-10029 |
| AF647 donkey anti-goat IgG | Invitrogen | A21447 |
| DAPI (1 mg/mL) | Thermo Scientific | 62248 |
| Fluoromount-G | Invitrogen | 00-4958-02 |
| 24x55mm No. 1.5 Thickness Cover Slips | Epredia | 152455 |
| ***PhenoCycler Antibody Conjugation and Tissue Staining*** |  |  |
| Lammeli Loading Dye | Bio-Rad | #1610737EDU |
| GelCode Blue Stain Reagent | Thermo Scientific | 24590 |
| 1X D-PBS | Wisent | 311-425-CL |
| Methanol | Commercial Alcohols | P016MEOH |
| ***Akoya Reagents*** |  |  |
| 10X PhenoCycler Buffer | Akoya | 7000001 |
| Staining Kit | Akoya | 7000008 |
| Conjugation Kit | Akoya | 7000009 |
| Black-walled 96-well plate | Akoya | 7000006 |
| Adhesive foil | Akoya | 7000007 |
| Assay Reagent | Akoya | 7000002 |
| Nuclear Stain | Akoya | 7000003 |
| Flow Cell | Akoya | 240204 |
| ***Software*** |  |  |
| QuPath | Bankhead et al (67) |  |
| StarDist | Schmidt et al (68) |  |
| MatLab | MathWorks |  |
| CytoMAP | Stoltzfus et al (53)<br>Weigert et al (69) |  |
| GraphPad Prism | Dotmatics |  |
| ***Other*** |  |  |
| Microtome | Leica | RM2125 RTS |
| PT Link for Pre-Treatment | Agilent |  |
| LED Lamps |  | 20000 Lux Intensity |
| AxioScan 7 | Zeiss |  |
| PhenoCycler-Fusion | Akoya |  |

### Antibody Conjugation to an Oligonucleotide Barcode

Once an antibody has shown strong and specific signal by IF, it can proceed to conjugation. Antibodies can be conjugated to barcodes which have complementary reporters in ATTO550, AF647, or AF750 fluorophores. IF screening will inform which fluorophore will give optimal results. In general, antibodies which show very strong and specific staining should be conjugated to barcodes that have complementary reporters in AF750, antibodies which have weaker signal and lower abundance should be conjugated with barcodes that have complementary reporters in in AF647, and antibodies which have medium abundance and weak to medium signal strength should be conjugated with barcodes that have complementary reporters in ATTO550.

Antibody conjugation requires reagents from Akoya Biosciences, and thus follows their recommended protocol. A more detailed protocol can be found here: https://www.akoyabio.com/wp-content/uploads/2021/01/CODEX-User-Manual.pdf

#### Pre-experiment Notes

- Antibodies to be conjugated <u>must be carrier-free</u>. The presence of BSA or other stabilizing agents will interfere with conjugation.
- If conjugating more than one antibody, carefully label all MWCO columns prior to starting. We recommend conjugating no more than 3 antibodies at a time, to reduce the risk of cross-contamination.
- Reagents which are purchased from Akoya and used “as-is” will be annotated as (*A*). Reagents that are purchased from Akoya but need preparation prior to use will be <u>underlined</u> and annotated as (*A*).

#### Conjugation Reaction

1. For each antibody to be conjugated, add 450 µL of Filter Blocking Solution (*A*) to a labelled 50 kDa MWCO column, then spin at 12,000g for 2 minutes. Following centrifugation, discard flowthrough and aspirate any remaining liquid out of the filter unit.

i. Note: This is the only step where the liquid should be aspirated out of the filter unit. In all subsequent steps, the remaining liquid contains the unconjugated/conjugated antibody.
2. Add 50 µg of each antibody to be conjugated to their respective filter units, at an adjusted volume of 100 µL. Spin down tubes at 12,000g for 8 minutes, and discard the flowthrough.
3. Add 260 µL of <u>Antibody Reduction Master Mix (*A*)</u> to the top of each filter unit, close this lid, vortex for 3 seconds, then allow to sit at room temperature for 30 minutes.

i. Note: do not allow this reaction to exceed 30 minutes, as it can result in irreversible antibody damage.
4. Spin down tubes at 12,000g for 8 minutes, then discard the flowthrough.
5. Add 450 µL of Conjugation Solution (*A*). Spin down again at 12,000g for 8 minutes, then discard the flowthrough.
6. During the second centrifugation, prepare each assigned Barcode (*A*) by adding 10 µL of molecular biology grade nuclease free water, then add 210 µL of Conjugation Solution (*A*) to the resuspended barcodes.
7. Add the barcode solution to the filter. Close the lid and vortex for 3 seconds. Incubate the antibody conjugation reaction at room temperature for 2 hours.
8. Spin down tubes at 12,000g for 8 minutes, then discard the flowthrough.
9. Add 450 µL of Purification Solution (*A*) to each filter, and spin down tubes at 12,000g for 8 minutes, then discard the flowthrough.
10. Repeat Step 9 for a total of 3 purifications. At the end of the third purification, the filter will contain the conjugated antibody.
11. For each antibody, label a fresh tube with the antibody name and the barcode ID. Add 100 µL of Antibody Storage Solution (*A*) to each filter. Then, invert the filter unit into the new collection tube, and spin down at 3,000g for 2 minutes.

i. Note: The final volume of the antibody will be around 120 µL
ii. Note: For long term storage, transfer antibodies to autoclaved screw top tubes, to reduce evaporation.

#### Validation of Conjugation to an Oligonucleotide Barcode

12. Cast a 10% SDS-PAGE gel, with 2 wells for each antibody whose conjugation is being validated, plus an additional well for the protein ladder (ie. if validating 4 antibodies, you would need a total of 9 wells, so a 10-well gel will suffice). Set up gel running apparatus, as you would for a typical western blot.

i. Note: Details on SDS-PAGE gel casting can be found here: https://www.bio-rad.com/webroot/web/pdf/lsr/literature/Bulletin_6201.pdf
13. Add 1 µL of unconjugated antibody to a tube with 9 µL of 1X lammeli loading dye. Add 0.5 µL of conjugated antibody to a different tube with 9.5 µL of 1X lammeli.
14. Boil samples for 5 minutes on a heating block at 95 °C.
15. Load samples and protein ladder into the gel and run until resolved.

i. Note: We typically run our gels at 90 V for 1.5 hours.
16. Following running, carefully remove the gel from the cassette, and place in a glass container. Fill the glass container with GelCode Blue Reagent.
17. Allow the gel to incubate in the GelCode reagent with gentle rocking, until the solution changes from pale brown to blue.
18. Carefully discard the GelCode reagent and replace with distilled water. Allow the gel to rinse with gentle rocking for 20 minutes. Wash 3 times with distilled water in the same fashion for 20 minutes each.
19. Following washing, blue antibody bands should resolve around 50 kDa. Image the bands with any gel imaging apparatus, such as a ChemiDoc.
20. Conjugation occurred successfully if there is an upward shift in weight from the unconjugated antibody to the conjugated antibody.

### Optimization of Conjugated Antibodies

Prior to performing a complete PhenoCycler experiment, conjugated antibodies must be further quality controlled and titrated. To do this, tissues are stained with the conjugated antibody of interest, and PhenoCycler reporters are manually applied and imaged. Staining fidelity is then assessed, and proper staining conditions are noted for larger multiplexed staining experiments.

#### Tissue Staining and Fixation

1. Follow steps 1 – 3 for *Deparaffinization and Antigen Retrieval*, described above.
2. To quench auto-fluorescence, place the slide in glass container and cover with <u>Bleaching</u> <u>Solution</u>. Sandwich the glass container between two LED lamps for 45 minutes at room temperature.
3. Replace the <u>Bleaching Solution</u> with fresh <u>Bleaching Solution</u> and repeat LED photobleaching for 45 minutes at room temperature (72).

i. Note: we find that this extended LED photobleaching step helps decrease auto-fluorescence associated with FFPE tissue staining.
ii. Note: The amount of H_2_O_2_ can be increased to 10% in tissue which demonstrate high levels of autofluorescence, such as heart or liver.
4. Wash the tissue 4 times in 1X PBS for 5 minutes per wash.
5. Dry the glass around the tissue with a Kimwipe, and circle tissue with a PAP pen.
6. Cover the tissue with Staining Buffer (*A*) and allow the tissue to equilibrate at room temperature for 30 minutes.
7. While the tissue is equilibrating, prepare the antibody solution. Antibodies are diluted in <u>Staining Buffer, completed with N Blocker, G2 Blocker, J Blocker, and S Blocker (*A*).</u>
8. Stain tissue by adding prepared antibody onto the tissue.

i. Note: Staining and time and temperature need to be optimized for each antibody. Common staining conditions include 3 hours at room temperature, or overnight at 4 °C.
9. Following antibody incubation, wash tissue 3 times in fresh Staining Buffer.

i. Note: For highly multiplexed experiments where antibody staining conditions differ, staining can be done sequentially. For instance, 3 antibodies can be applied for 30 minutes at 37 °C, then tissue can be washed in buffer and the remaining antibodies in the staining panel can be applied overnight at 4 °C.
10. Perform first tissue fixation, by incubating tissue in <u>Post-Staining Fixation Solution (*A*)</u> for 10 minutes at room temperature. Rinse tissue 3 times with PBS.
11. For the second fixation, transfer slides to a Coplin jar on ice filled with pre-chilled methanol. Allow to incubate for 5 minutes, then quickly transfer back to PBS. Rinse 3 times with PBS.
12. For the third and final fixation, add <u>Final Fixative Solution (*A*)</u> to slides, and incubate in a humidity chamber at room temperature for 20 minutes. Rinse tissue 3 times with PBS.
13. Transfer slide to Coplin jar with Storage Buffer (*A*).

i. Note: Slides can remain in Storage Buffer (*A*) at this step for up to 5 days at 4 °C.

#### Manual Application of PhenoCycler Reporters and Tissue Imaging

14. Prepare <u>Screening Buffer (*A*)</u> and allow to equilibrate to room temperature for 20 minutes before use.
15. Rinse slides in 3 changes of <u>Screening Buffer (*A*)</u> for 1 minute each, to allow the tissue to equilibrate to the new buffer.
16. Prepare the <u>Reporter Stock Solution (*A*)</u> and add 2.5 µL of each reporter to be tested to 97.5 µL of <u>Reporter Stock Solution (*A*).</u>

i. Note: More than one antibody/reporter can be tested at a time, provided the reporters are conjugated to different fluorophores. For instance, if tissue is stained with CD4-BX001 and CD19-BX002, 2.5 µL of both RX001-AF750 and RX002-ATTO550 can be diluted into 95 µL of <u>Reporter Stock Solution (*A*)</u> for marker visualization in a single step.
17. Pipette the prepared <u>Reporter Stock Solution (*A*)</u> onto the tissue and incubate protected from light for 5 minutes.
18. Rinse slides in 3 changes of <u>Screening Buffer (*A*),</u> for 1 minute each.
19. Rinse slides with 1 change of <u>1X PhenoCycler Buffer (*A*).</u>
20. Mount coverslips onto slides with Flouromount-G, and then allow to dry for 15 minutes.
21. Image slides with the same microscope that will be used for PhenoCycler image acquisition.

#### Assessing PhenoCycler Staining

When assessing the quality of a conjugated antibody, it is important to keep in mind the SNR from the previous step, as it will be used as a reference to compare for quality control. At this stage, multiple antibody concentrations should be tested as well as multiple incubation times and temperatures in order to get the best SNR. We also recommend performing one final staining with two extra markers: one that should co-localize and one that should not with the conjugated antibody being tested. This step will allow you to assess any non-specific binding of conjugated antibody and adjust staining and acquisition parameters for best SNR. Staining intensity and pattern should match the one obtained by standard IF staining.

### PhenoCycler Multiplexed Imaging

Once all antibodies have been conjugated and optimized, you may proceed to a full PhenoCycler staining experiment. Prior to beginning, all antibodies must be assigned to a cycle, a step that requires some thoughtful consideration. Each cycle will consist of up to 3 different antibodies, conjugated to barcodes that have reporters with different fluorophores. For instance, cycle 2 may consist of imaging CD4-BX001, CD19-BX002, and CD11b-BX003, which have RX001-AF750, RX002-ATTO550, and RX003-AF647 complementary reporters. When designing cycles, we try to include markers that are not likely to be present on the same cell type (ie, CD4, a marker of helper T cells, may be put in cycle 2, while CD3, a pan-lymphocyte marker, may be put in cycle 3). The first and last cycle of each staining experiment will consist of only DAPI (“Blank” cycle).

#### Tissue Staining and Reporter Plate Preparation

1. Follow steps 1 – 13 for *Tissue Staining and Fixation*, using all conjugated antibodies in the staining panel. Leave slide in Storage Buffer (*A*) until prepared to proceed to a full PhenoCycler Image Acquisition run.

i. Note: For full PhenoCycler staining experiments, antibodies should not exceed 40% of the total Complete Staining Buffer solution, or insufficient blocking will occur.
2. Prepare enough <u>Reporter Stock Solution (*A*)</u> for the number of cycles in the experiment (each cycle requires a maximum of 250 µL of <u>Reporter Stock Solution (*A*</u>)).
3. For each cycle, label an amber 1.5 mL Eppendorf tube, and add 5 µL of each reporter for the assigned cycle. Complete to a volume of 250 µL using <u>Reporter Stock Solution (*A*)</u>. Mix the contents gently by pipetting up and down.

i. Note: keep reporters on ice until use, and spin down prior to pipetting to collect any accumulated liquid in the cap.
ii. Note: the first cycle and the final cycle will consist of <u>Reporter Stock Solution (*A*),</u> with no florescent reporters added (ie. “Blank” cycles)
4. For each assigned cycle, pipette the reporter solution into a black-walled 96-well plate. Cover the wells with adhesive foil.
5. The reporter plate can be stored at 4 °C for up to two weeks or can be used immediately for the PhenoCycler experiment.

#### PhenoCycler Image Acquisition

Images are acquired using the default PhenoCycler protocol. In this study, we used the Phenocycler-Fusion system combining Phenocycler instrument with the Fusion microscope to streamline acquisition. We used acquisition parameters of the different antibodies defined during the titration step to acquire the fully stained tissue.

### Open-Source Data Analysis

Following a complete PhenoCycler staining experiment, PhenoCycler software will process images for downstream analysis. Imaging processing includes tile stitching and background correction. The final multiplexed image will be in QPTIFF format, which can be imported and visualized by many image analysis programs.

In this pipeline, we use the open source QuPath software, v3.2, which can be found here: https://github.com/qupath/qupath/releases/

Cell segmentation is achieved using StarDist, which can be found here: https://github.com/qupath/qupath-extension-stardist/releases

The pre-trained model we used for StarDist Segmentation can be found here: https://github.com/qupath/models/tree/main/stardist

The StarDist .groovy file used in this study and sample Classifier data can be found here: https://github.com/MMdR-lab/mouseCODEX-paper

#### Setup

1. Create directory including StarDist segmentation extension (qupath-extension-stardist-0.4.0.jar), the pre-trained StarDist model (dsb2018_heavy_augment.pb), and stardist_segmentation_0.5px.groovy file.
2. Set shared script directory with the command Automate -> Shared scripts -> set script directory and select directory including .groovy file and .pb file.
3. Create a list of the channel names in the order of acquisition in a .txt file with a separate line for each name.

#### QuPath Image Import

1. Create a new project in QuPath and add the PhenoCycler QPTIFF as a new image. Double click to open the image, and a pop-up will prompt you to select the image type. Set the image type as Fluorescence and keep “Auto-generate pyramids” selected.

i. Note: QPTIFF files are generated by the Phenocycler-Fusion system. For researchers using the Phenocycler combined to standard microscope, single channel OME-TIFF files can be combined into multiple channel OME-TIFF in ImageJ prior to proceeding.
2. Once the QPTIFF image is opened, all markers (i.e. αSMA, CD3, CD4, CD8, CD11b, CD11c, CD19. CD31, CD45, c-Myc, F4/80, FoxP3, Ki67, MelanA, MPO, and NaKATPase) will be simultaneously visible on the tissue, labeled as the fluorophore they were conjugated to in the order of cycle acquisition.
3. To set channel names, copy the list of channel names to the clipboard and then select the corresponding channels in the “Brightness/Contrast” dialog from the “View” dropdown menu and paste. Click apply to confirm.
4. In the “Brightness/Contrast” dialog box, you can toggle markers on and off, change their pseudo-colouring, and adjust their min/max display.
5. Make the channel names available as classifications in the “Annotations” tab by right - clicking or selecting the vertical ellipsis next to “Auto set” and choosing “Populate from image channels”.

#### QuPath Cell Classification and Cell Segmentation

6. To classify cells into phenotypes, a training image is used. The training image will contain 5 or 6 representation regions of interest, pooled into a single image.

i. To create a training image, select “Training images” from the “Classify” dropdown menu, and select “Create region annotations”.
ii. Using the default settings of: Width-500; Height-500; Size units-µm; Classification-Region*; and Location-Viewer Centre, create regions throughout the tissue which contain the cell phenotypes you wish to annotate.
iii. Save the image.
iv. From the “Classify” dropdown menu, select “Training images”, and select “Create training image”.
v. From the popup menu, select “Region*” as the Classification, type “50,000” px as Preferred image width, and toggle “Rectangles only”, then click OK.
vi. A training image will appear in the Project Image List dropdown menu.
vii. Open the training image, and save the project.
7. To segment the training image into cells, StarDist is used. Using the rectangle annotation tool, select the entire region.
8. To segment the annotated region into cells, select StarDist Cell Segmentation from the shared scripts in the Automate dropdown. QuPath. When the script editor appears, select “Run”.

i. Note: If an annotation is not selected, the error “Please select a parent object!” will appear.
9. A dialog box will appear, prompting the selection of the segmentation file. Choose the dsb2018_heavy_augment.pb file located in the directory you created in step 1.
10. Once segmentation is complete, you will be able to see cell detections in red overlay on the image. You can toggle the visibility of the cell detections using the overlay capacity slider bar at the top of the image window.
11. To proceed with cell classification, from the “Classify” dropdown menu, select “Training images” and select “Create duplicate channel training images”. From the popup window, select the markers that you wish to use to enable cell classification. Check the “Initialize Points annotations” box then select “OK”. There will now be duplicate training images in the Project Image List dropdown menu for each marker in the staining panel. These duplicate channels will be used for manual annotation of cell phenotypes.

i. Note: cell classification should be done in a single duplicate training image for phenotypes that are characterized by mutually exclusive markers (ie. lineage markers). For instance, if CD8+ T cells are classified as CD3+ CD8+, macrophages are classified as CD11b+ F4/80+, and fibroblasts are classified as CD45-αSMA+, they can be used in a classifier together.
ii. Note: in this project, we trained two classifiers to detect a total of 10 cell types. The first classifier was trained to detect CD8, FoxP3, CD31, F4/80, and CD11c. The second classifier was trained to detect CD4, CD19, MPO, αSMA, and Ki67+ tumour cells.
12. Open the duplicate image for the first cell type(s) you wish to classify.
13. Open the points annotation tool, add an annotation, and right click to set the annotation class (ex. if you are classifying helper T cell, set the class to CD4). Add a second annotation and set the class to “Ignore*”.
14. Using point annotation, annotate 30-60 cells of your class of interest, and annotate another 30-60 cells as “Ignore*”. The “Ignore*” cells should be mutually exclusive from the cell you are classifying. For instance, if you are classifying CD4+ T cells, you could select CD8+ T cells, B cells, or tumour cells for the “Ignore*” class. This helps train the classifier to better detect your cells of interest.
15. From the “Classify” dropdown menu, select “Train object classifier”.

i. Set Object filter to “Cells”
ii. Set Classifier to “Artificial neural network (ANN_MLP)
iii. Set Feature to “All measurements”
iv. Set Classes to “Selected classes”
v. Set Training to “Unlocked annotations”
16. Click “Live update”. The cell mask on the training image should update to show where your cell phenotype has been detected.
17. Manually assess if the cell classifier is accurately detecting your cell phenotype of interest. If there are many false positive detections, continue to add annotations for “Ignore*”. If there are many false negative detections, continue to add annotations for your cell type of interest.
18. Once you are content with the cell classifier, enter the object classifier name, and click “Save”.
19. Repeat steps 12 – 18 for all cell phenotype classes you wish to annotate in your tissues.
20. Open the main image from the Project Image List.
21. Using the rectangle or polygon annotation tool, select the regions you wish to analyze. Following steps 7 – 9, use StarDist to segment the annotation region into cells.
22. Classify cells into phenotypes by opening the “Classify” dropdown menu, selecting “Object classification”, then selecting “Load object classification”. Select the classifiers you wish to apply to the tissue, then select “Apply classifiers sequentially”.

i. Note: If more than one classifier is used to detect cell types, there may be redundancy in classification (ie, some cells will be annotated as more than one class). For instance, in this study, our first classifier detected FoxP3+ cells, and our second classifier detected CD4+ cells. Thus, when the classifiers were applied together, regulatory T cells were classified as FoxP3+ CD4+.
ii. Note: Due to cell segmentation noise, sometimes dual classifiers may not make biological sense. It is up to the researcher to manually assess each cell class, and collapse classes as necessary.
23. Now, each cell will be annotated as a Phenotype. To export this data for spatial analysis with CytoMAP:

i. Save the QuPath project.
ii. From the “Measure” dropdown menu, select “Export Measurements.”
iii. Select the image you wish to export measurements from, and choose “cells” as the export type. Change separator type to “Comma (.csv)”.
iv. Click “Populate”, then select the columns to include from the dropdown list: Image Name, Image, Class, Centroid X, Centroid Y, and Cell Mean for each marker. The resulting .csv file will contain the fluorescence intensity of each marker for each cell within the image, plus all cells will be annotated for their cellular phenotypes.

#### CytoMAP Spatial Analysis

24. In MATLAB, install the CytoMAP plugin in the “Add-Ons” drop down menu.

i. For desktop use without MATLAB downloaded, an installer for the compiled version of CytoMAP is available at https://gitlab.com/gernerlab/cytomap/-/tree/master/. Follow the installation prompts
25. Open CytoMAP. From the “File” dropdown menu, select “Load Table of Cells”, then select the .csv file generated in step 23.

i. Be mindful of .csv formatting when uploading. CytoMAP may not recognize certain symbols, such as ampersands or slashes.
26. A popup dialog box will prompt you to select the X axis. Click “Ch_Centroid_X_m” and click “Okay”.
27. A popup dialog box will prompt you to select the Y axis. Click “Ch_Centroid_Y_m” and click “Okay”.
28. A popup dialog box will prompt you to select the Z axis. Click “There is no Z (make a fake one)” and click “Okay”.
29. A “File Import Options” box will pop up. Select “Load”.
30. Select “Annotate Clusters”, and from the Select Classification Chanel dialog box, select “Ch_Class”. From the Annotate Class popup box, select “Save Annotations”. Close the Save Annotations box.
31. To make a heatmap showing the cellular mean intensity of the markers in the staining panel within the different cell phenotypes, click the “Extensions” dropdown menu, and select “cell_heatmaps.m”

i. Choose the cell phenotypes you wish to include.
ii. Choose the Channel MFIs you wish to include.
iii. Normalize per Sample.
iv. Select “MFI normalized to mean MFI of all cells”.
v. Select “Phenotype” for what to compare.
vi. Select “Individual Heatmap for each Sample”
vii. Select “linear” for scale.
viii. Click “Okay”.

i. Note: If multiple .csv files are imported and annotated, you may choose to generate a combined heatmap.
32. To cluster cells into neighborhoods, select “Define Neighborhoods”.

i. Choose “Raster Scanned Neighborhood” for Neighborhood Type.
ii. Type “50” for Neighborhood Radius.
iii. Select “Fast Way”
iv. Click “Okay”
33. Once the loading bar for Defining Neighborhoods has finished, click “Cluster Neighborhoods into Regions”.

i. Select all Phenotypes for sorting.
ii. Use setting “Composition: Number of Cells/ Number of Cells in Neighborhood”
iii. Use setting “MFI normalized to mean MFI per neighborhood” and Normalize per Sample.
iv. For Colour scheme, select “sum(y,2)
v. For Number of Regions, select “Davies Bouldin (default)”
vi. For Model name, select “Create New Model”
vii. For Data Input Type, select “Raster Scanned Neighborhood”.
viii. For Algorithm, select “NN Self Organizing Map”.
ix. Click “Okay”
x. Enter a unique name for the Model.
34. Two figures will popup, one showing the Number of Clusters and the Davies Bouldin values, and the other showing the newly defined regions superimposed on the tissue image.

i. Note: In tumour tissues, overall cellular disorganization leads to fewer definitive regions.
35. To generate a heatmap showing the spatial relationships between cells in the tissues, select “Cell-Cell Correlation”.

i. Select the Phenotypes you wish to include.
ii. For Neighborhood Type, select your unique Model name.
iii. For data preparation, select “Cellularity: Number of Cells / Neighborhood”.
iv. Normalize per Sample.
v. Select “Individual Heatmap for each Sample”.
vi. For Colour Scale, select “linear”.
vii. For Calculation, select “Pearson Correlation Coefficient”.
viii. For Transform, select “None”.
ix. For Confidence Interval, select “1”.
36. CytoMAP can be used for other types of spatial analysis, and details can be found here: https://cstoltzfus.com/posts/2021/06/CytoMAP%20Demo/

## Author Contributions

Conceptualization – M.J.A., C.G., K.K.M., S.V.d.R.

Methodology – M.J.A., C.G., P.M., V.G., H.C.

Investigation – M.J.A., C.G., V.G., H.C., S.E.J.P., F.H.

Animal Models – M.J.A., S.E.J.P., F.H., N.G.

Data Visualization – M.J.A., C.G., P.M.

Supervision – N.J., W.M., K.K.M., S.V.d.R.

Writing – Original Draft – M.J.A

Writing – Reviewing and Editing – all authors

